# Identification of *C. elegans* strains using a fully convolutional neural network on behavioural dynamics

**DOI:** 10.1101/433052

**Authors:** Avelino Javer, André E.X. Brown, Iasonas Kokkinos, Jens Rittscher

## Abstract

The nematode *C. elegans* is a promising model organism to understand the genetic basis of behaviour due to its anatomical simplicity. In this work, we present a deep learning model capable of discerning genetically diverse strains based only on their recorded spontaneous activity, and explore how its performance changes as different embeddings are used as input. The model outperforms hand-crafted features on strain classification when trained directly on time series of worm postures.

## 1 Introduction

Animals interact with the world through their behaviour which involves the processing of sensory inputs and the generation of motor outputs by the neural system. Until recently most of the studies of animal behaviour relied on manual scoring certain expert defined actions by human reviewers. While time consuming, these approaches also lack objectivity and can result in poor reproducibility of experimental results. Advances in recording, storage and processing technology make it now possible to collect time-lapse recordings, and analyse large data collections in a controlled manner[1]. Inspired by the recent advancements in recognising human actions [15, 14, 3], computer vision methods are now being adopted to develop computer assisted approaches for the quantitative analysis of animal behaviour [9, 8, 17, 6].

The nematode worm *C. elegans* is particularly appropriate for behaviour quantification. Due to its experimental amenability and its small nervous system, it is the perfect candidate to understand the genetic basis of behaviour[17] and to dissect the neural circuits responsible for complex behaviours such as foraging, navigating and mating[6]. Even more it has a simple morphology that can be abstracted as the coordinates of its midline (skeleton). This abstraction has been shown to be effective in characterising worm behaviour particularly in large sets of data [19, 20, 2, 16, 21, 12].

Recently proposed convolutional networks for object recognition [18, 10] and semantic segmentation [13] not only demonstrate that such approaches can out-perform traditional methods, they also illustrate that it is possible to learn such models directly from raw image data. Here, we aim to investigate if biologically relevant motion signatures can be learned directly from video data. To achieve this goal we developed a framework for training a deep learning classifier that can predict the worm’s strain type based on its recorded behaviour. We consider diverse sets of inputs for the model: some derived from the worm skeletons extracted using traditional computer vision methods, while others are learned using an autoencoder on the raw images.

The datasets used in our study are described in Section 2. The different approaches for extracting relevant signatures that capture the pose of the worm are detailed in Section 3. The deep learning based model for classification is presented in Section 4. Finally, we present our results in Section 5 and summarise our conclusions. Overall our results are very promising. When using inputs that are derived directly from the worm postures the classifier outperforms the accuracy obtained using hand-crafted features. However, although the image reconstruction results obtained by the autoencoder are very convincing, the classification performance degrades when the autoencoder embeddings are used as inputs. Potential approaches to improve the classification accuracy are discussed in Section 6.

## 2 Datasets

All the videos were segmented, tracked, and skeletonised using Tierpsy Tracker[7] (http://ver228.github.io/tierpsy-tracker/).

### Single-Worm (SW) Dataset

The data was obtained from the Open Worm Movement Database[7](http://movement.openworm.org/). Each of the videos in this dataset focuses on a single worm that is followed, with the help of a motorised stage, as it moves around the recording plate. We restricted our analysis to include only 15 minutes videos of young adults on food, and where at least 50% of the frames were successfully skeletonized by the tracking software. We include mostly videos that were used in published papers[2, 16].

The dataset comprises 10476 videos of individual worms divided between 365 different classes. It includes mutants of the laboratory control strain (N2) affecting neurodevelopment, synaptic and extrasynaptic signalling, muscle function, and morphology as well as wild isolates representing some of the natural heterogeneity of *C. elegans*. All the videos for a given strain are given the same class label, except N2 where hermaphrodites and males are are considered as separated classes. We only included classes that have at least six videos.

### Multi-Worm (MW) Dataset

The dataset consists of recordings of a fixed 2×2mm area taken with a high-resolution camera. There are either 5 or 10 worms in each of the recorded plates. The twelve strains used in this dataset are part of the divergent set from the Caenorhabditis elegans Natural Diversity Resource (CeNDR)[4]. This small subset of CeNDR collection is a representative sample of the genetic diversity found among the *C. elegans* wild isolates. A total of 308 videos were collected, with between 25 and 28 videos per strain.

One important difference with the SW dataset is that due to the overlap between worms trajectories the identity of individual worm is frequently lost. The results is that rather than having continuous data for each worm along the video, we have a series of fragments terminated every time two or more worms encounter.

## 3 Postural Embeddings

One key consideration in our study is how the pose of a given worm should be embedded to facilitate the classification task. The skeletonization summarises the worm posture and its head to tail orientation. Therefore embeddings extracted from skeletons have the advantage that they implicitly take the anatomy of the worm into account. Alternatively, it is possible to learn a representation directly from the image data using an autoencoder. In the following sections we provide the details for each of these approaches. The embeddings are then stacked over time to create the postural maps presented in Fig. 1 and fed to the classifier as explained in the next section.

**Fig. 1.**
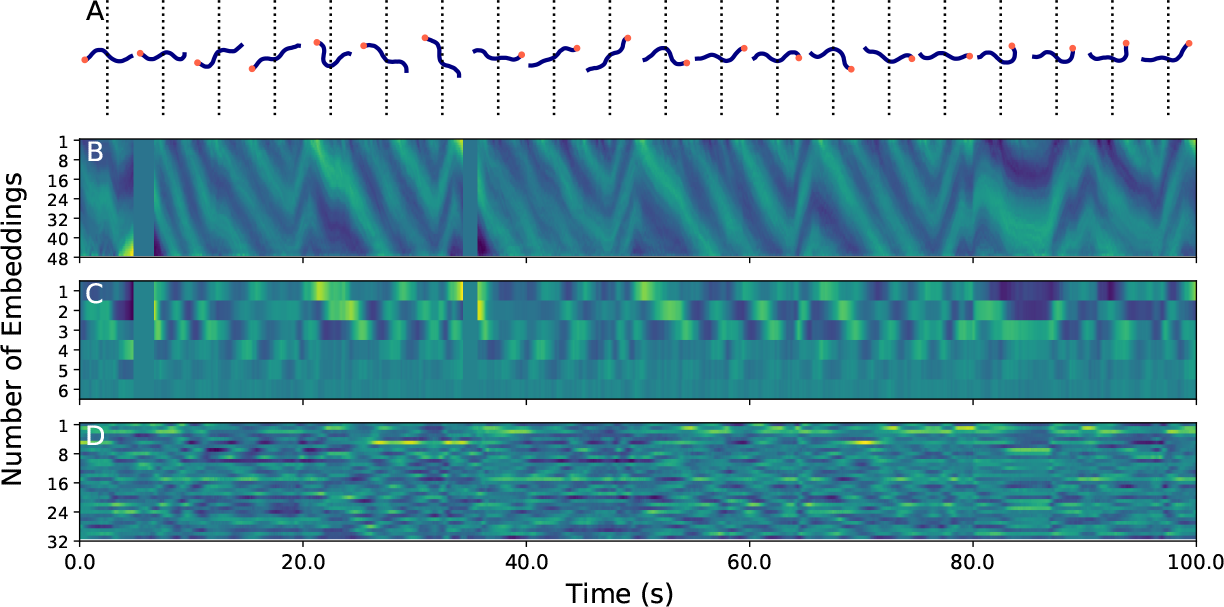
Comparison between the different types of embeddings for the same worm movie. (A) Example of the worm postures at intervals of 128 frames. The blue lines represent the worm skeletons, the orange circles show the head side and the dash lines align to the corresponding column in the maps below. (B) Skeletons angles, the elements in the embeddings are ordered from head to tail. (C) Eigenworms, the elements are sorted from the most important PCA to the least important. (D) Embeddings from the autoencoder.

### 3.1 Skeletons Angles and Eigenworms

As a preprocessing step the skeletons are interpolated in space to have a total of 49 evenly spaced points. The skeletons are interpolated also in time to achieve a constant temporal sampling separation of 0.04*s*. Due to clutter, artefacts or the worm coiling over itself the skeletonisation can fail. If the time gap of unskeletonised frames is less than 0.25*s* linear interpolation between the closest skeletonised frames is used to compensate for the missing data. Larger gaps coming mostly from coiling/turning worms are set to zero. Finally the skeletons are smoothed in both space and time using the Savitzky-Golay filter.

The dimensions of each skeleton is 49×2. To further reduce the dimensionality and focus on the posture, we follow the procedure introduced in ref[19]. As a first step we calculate the tangent angle between consecutive points as

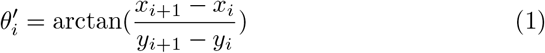

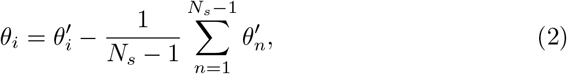

where *N*_*s*_ = 49 is the total number of segments, *x*_*i*_ and *y*_*i*_ are each segment coordinates, and *θ*_*i*_ is the corresponding segment angle. The resulting embedding has 48 elements as shown in Fig. 1B.

In a second step the angles are projected onto a set of eigenvectors *u*_*μ*_ previously calculated from the PCA of all the skeletons angles in the SW dataset, also called eigenworms. Stephen *et al.*[19] demonstrated that the first four eigenvectors are sufficient to capture 95% of the observed postural variance. Following Li *et al.*[12] we decided to use the first six eigenvectors that capture 98% of the total variance. An example of the resulting embeddings is shown in Fig. 1C.

### 3.2 Autoencoder

For the MW dataset we also extracted the embeddings directly from videos using the convolutional autoencoder described in Fig. 2A. Since most of the pixels in the video are background we only use regions of 128×128 pixels around each individual worms. We set the embedding dimension to have 32 elements. We used 95% of the videos for training and the rest was reserved as the test set. During training we used data augmentation by applying random shifts and flips to the images. We used the *L*_1_-norm as loss function, and train with stochastic gradient descent (SGD) using a learning rate (lr) of 10^−4^ and momentum of 0.9. We stopped the training when the loss function in the test set did not show any improvement. Some examples of the encoded/decoded images are shown in Fig.2B.

**Fig. 2.**
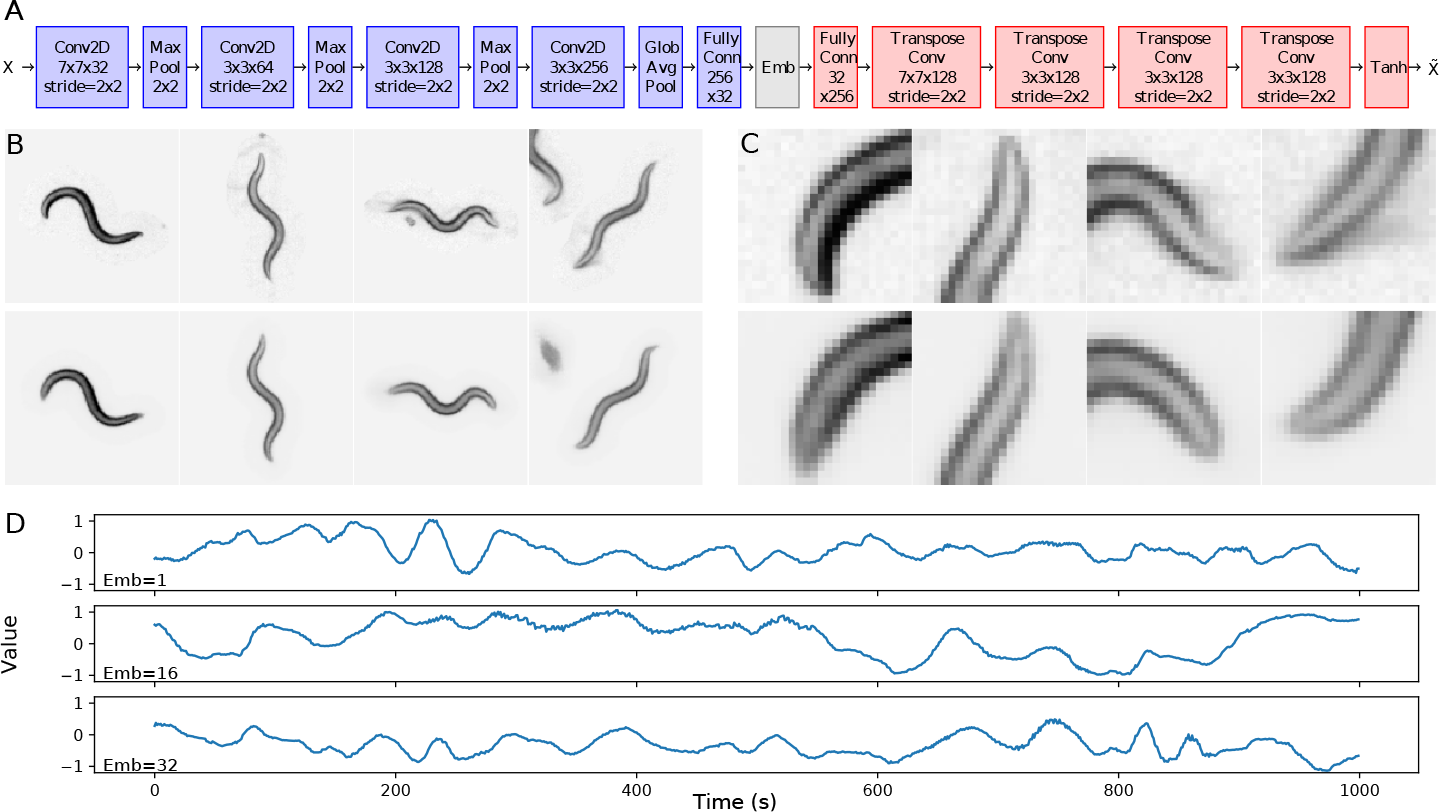
The embeddings learned by the autoencoder can recover the worm shape with high accuracy. (A) Autoencoder architecture. The blue blocks show the encoder module, the grey block the embedding, and the red blocks the decoder module. (B) Comparison between the original image, on top, and the autoencoder reconstruction, bottom, of four randomly selected images. The reconstruction is highly similar to the original and it even suppresses objects surrounding the worm in the centre of the image. It is likely that this denoising behaviour arises from the model regression to the sample mean[11]. (C) Same as B but zoomed on the worm head. (D) Example of three elements of the embedding vector over time. The signals are continuous indicating a smooth temporal transition between the embeddings for different postures in the video.

## Classification

The classification model is shown in Fig. 3 and was inspired by the Dilated Resnet Network architecture [23]. Contrary to most classifiers with stacked convolutions, this architecture does not aggressively reduce the network output size. The result is that the output layers before the classifier generate feature maps rich in spatial information and capable of improve weakly-supervised tasks such as object location. These features maps could be interpreted as a timeseries feature transformation that together with the final global average is not different in spirit from the averages of user defined features used in Yemini *et al.* [22]. In brief, the model starts with a 7 × 7 convolution, followed by five 3 × 3 strided convolutions, then a series of dilated convolutions (2-4-2-1), and finally a classification layer consisting on global average pooling and a 1 × 1 convolution. Each convolution is followed by batch normalisation and a leaky ReLU activation. The strided convolutions modules condense the time dimension by a factor of 32 while the dilation layers add an extra factor of four. Therefore each row in the resulting features map contains the information of 128 frames. The statistics of the feature maps are then summarised by the global average pooling layer. One of the advantages of this model is that it can be applied to inputs of arbitrary size. We can use the same model with trajectories of different length or even combine several trajectories together in order to get predictions at the population level. This is particularly useful in the case of the MW dataset where the worm trajectories are fragmented and their individual identity is lost.

**Fig. 3.**
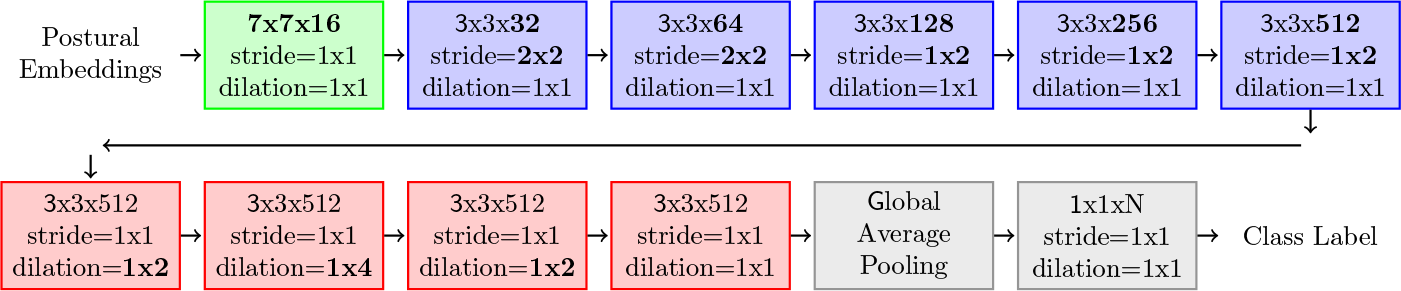
Architecture of the fully convolutional classification model. The model starts with a simple 7 × 7 convolution and is followed by a series of strided convolution. Since the time dimension in the embedding maps is much larger than the embeddings dimension, we used asymmetric strides 1 × 2 after the second strided convolution. We then used a series of dilated convolutions in order to increase the explored space without reducing temporal resolution as in ref[23]. Note that the dilation only occurs in the time dimension. Finally, we condensed the learned features using a global average pooling and create a classification layer using a 1 × 1 convolution.

### Training

We trained both datasets using SGD with a mini-batch size of eight. For the MW dataset model we used Adam as the optimiser with a lr of 10^−4^ for the angles and eigenworms, and a lr of 10^−5^ for the autoencoder embeddings. 75% of the data is used for training and 25% for testing. For the SW dataset we used a lr 10^−3^, a momentum of 0.9 and weight decay of 10^−4^. The lr is reduced to 10^−4^ when the loss reaches a plateau. Two thirds of the data are used for training and the rest for testing. In order to compensate for class imbalance, particularly problematic in the SW dataset, during training we sampled in two steps: first randomly selecting a strain, and then randomly selecting a video of that strain. Additionally, as a form of data augmentation, during training we concatenated the embeddings along randomly selected chunks of different trajectories until we completed a map with 22500 elements. During testing we evaluated the results concatenating all the embedding available for a given video.

### Comparison with manually-defined features

In order to have a baseline for comparison, we trained a simple classifier using the manually-defined features used in Yemini *et. al*[22]. This set of features is the best method reported in the literature to identify mutant strains from the control strain N2. The trained classifier consists in a fully connected layer followed by a softmax layer. Before training we z-transform data (subtracting the mean and dividing by the standard deviation) and set to zero any remaining undefined value. We trained using SGD with a lr of 10^−3^, a momentum of 0.9 and a mini-batch size of 64. We used the same data partition between training and test sets used for the fully convolutional classifier.

## 5 Results

### Classification accuracy

The classification results are presented in Table 1. Overall, the best results are obtained when the skeleton angles are used as input. A similar result is obtained in the MW dataset when the input of the model are the eigenworms, however for the SW dataset the top1 accuracy is almost ten percentage points lower. This suggests that there is some relevant information that is lost in the eigenworm transform. Nevertheless the eigenworms have the advantage of requiring less operations due to the smaller embedding size (6 vs 48) and by consequence the models can be trained considerably faster. This, combined with the fact that the eigenworms still produce better results than the manually-defined features makes them a promising alternative for small datasets. By contrast, when the autoencoder embeddings (Section 3.2) are used as input, the performance degrades significantly. It is hard to pinpoint what causes the drop in accuracy. The autoencoder produces denoised reconstructions of the original images. Therefore, the lower accuracy is not likely to be caused by a limited model capacity. One possible explanation might be that the embedding representation entangles the worm postures with other information presented in the images such as the texture or the worm orientation. This extra information might cause the model to overfit since it might be learning to identify individual videos rather than the actual differences between strains. One possible solution could be to condition the autoencoder embeddings to the worm skeletons. We plan to explore this possibility in detail in a future work.

**Table 1.** Classifier results.

| Training | Top-1 Acc. | Top-5 Acc | F1-score |
| --- | --- | --- | --- |
| MW Manually-defined features | 83.49 | 99.08 | 0.8323 |
| MW eigenworms | 98.17 | 99.08 | 0.9816 |
| MW angles | **98.17** | **99.08** | **0.9803** |
| MW autoencoder | 58.72 | 93.58 | 0.5944 |
| SW Manually-defined features | 43.20 | 66.47 | 0.3574 |
| SW eigenworms | 49.40 | 72.73 | 0.4204 |
| SW angles | **58.44** | **81.99** | **0.5323** |

### Learned features

We visualise the features learned by the classifier using t-SNE (Fig. 4). For the MW dataset the features cluster nicely along the strain type as it is expected from the model high classification accuracy. The SW data shows a larger overlap, particularly the control strain N2. This is not particularly surprising since most of the strains present in the database are mutants of N2 and some might not even have a clear different behavioural phenotype with respect to their parent strain. More interesting is to observe a clear cluster formed by the wild type isolates, as well as a degree of separation among the Uncordinate (Unc) and Egg-laying defective (Egl) strains[5].

**Fig. 4.**
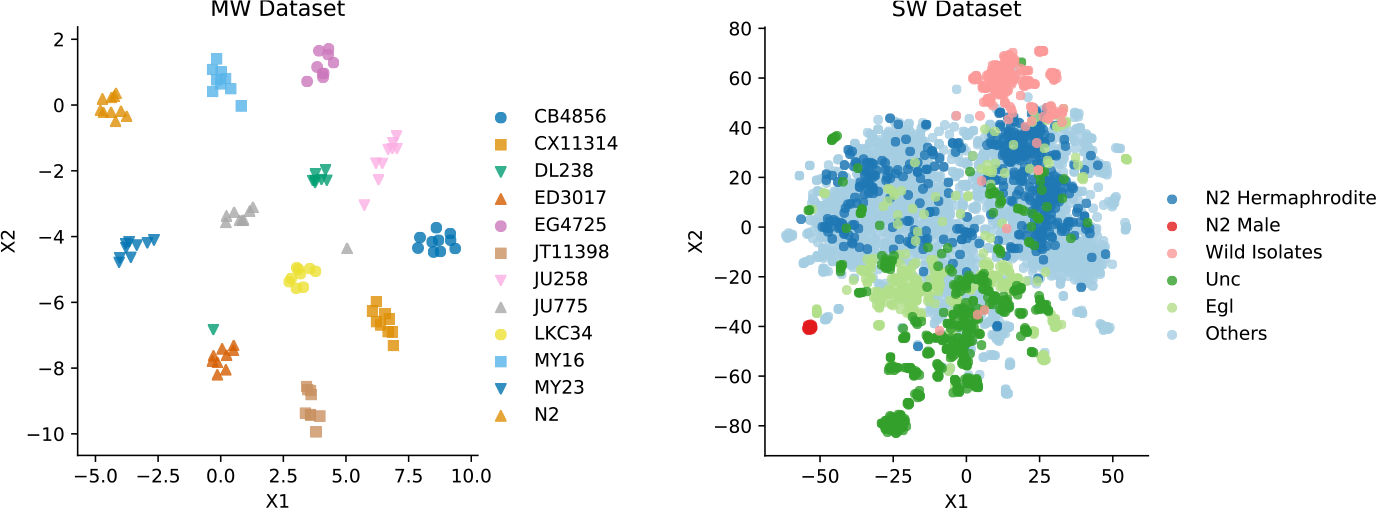
The features learned by the classifier cluster according to the strain type. t-SNE visualization of the activations of classifier penultimate layer on the test set data. (A) The features from the MW Dataset cluster tightly according to the strain type. The only exception is a DL238 video that clusters with ED3017 and it is likely to be a mislabelled sample. (B) The features from the SW Dataset show a higher degree of overlapping. For visualization purposes we grouped some strains that share similar behaviour. N2 hermaprodites, the control strain with 483 videos, N2 male with 19 videos, the wild isolates with 208 videos among 20 different strains, Unc, Uncoodinate meaning animals with deviations in self-propelled movement with 519 videos among 58 strains, Egl, Egg-laying defective with 357 videos among 38 strains.

## Conclusions

We have demonstrated that it is possible to train a classifier to distinguish between *C. elegans* strains using individual postural dynamics alone. More importantly, this classifier considerably improves the accuracy over the state of the art classification method given by the manual-crafted features defined in Yemeni *et. al* [22].

The main limitation of our current setup seems to be data overfitting rather than model capacity since all the trained models were able to fit almost perfectly their corresponding training data. More data should be available as the Open Worm Movement Database grows. Additionally, it might be possible to develop better sampling methods that could be used as data augmentation and help to reduce the early overfitting.

A logical next step will be to train an end-to-end model capable of classifying strains directly from the raw videos. However, to gather a dataset that contains the range of variability observed across laboratories remains a challenge. For example the SW dataset is probably the world largest worm behavioural dataset but all the data comes from a single laboratory using one type of setup. By contrast, there are several worm trackers available capable of extracting the skeletons from raw video. Those skeletons should not strongly depend on the imaging setup, and therefore a model trained on these inputs should be more easily deployed among the worm community.

## Acknowledgments

This work was supported by grants EPSRC SeeBiByte Programme EP/M013774/1 to JR and Medical Research Council MC-A658-5TY30 to AEXB. AJ benefited from both grants.

